# Proportionality: a valid alternative to correlation for relative data

**DOI:** 10.1101/008417

**Authors:** David Lovell, Vera Pawlowsky-Glahn, Juan José Egozcue, Samuel Marguerat, Jürg Bähler

## Abstract

In the life sciences, many measurement methods yield only the relative abundances of different components in a sample. With such relative—or *compositional—*data, differential expression needs careful interpretation, and correlation—a statistical workhorse for analyzing pairwise relationships—is an inappropriate measure of association. Using yeast gene expression data we show how correlation can be misleading and present *proportionality* as a valid alternative for relative data. We show how the strength of proportionality between two variables can be meaningfully and interpretably described by a new statistic *Φ* which can be used instead of correlation as the basis of familiar analyses and visualization methods, including co-expression networks and clustered heatmaps.

While the main aim of this study is to present proportionality as a means to analyse relative data, it also raises intriguing questions about the molecular mechanisms underlying the proportional regulation of a range of yeast genes.

## Author Summary

Relative abundance data is common in the life sciences; but appreciation that it needs special analysis and interpretation is scarce. Correlation is a popular as a statistical measure of pairwise association but should not be used on data that carry only relative information. Using timecourse yeast gene expression data, we show how correlation of relative abundances can lead to conclusions opposite to those drawn from absolute abundances, and that its value changes when different components are included in the analysis.

Once all absolute information has been removed, only a subset of those associations will reliably endure in the remaining relative data, specifically, associations where pairs of values behave proportionally across observations. We propose a new statistic *Φ* to describe the strength of proportionality between two variables and demonstrate how it can be straightforwardly used instead of correlation as the basis of familiar analyses and visualization methods.

## Introduction

Relative abundance measurements are common in molecular biology: nucleic acids typically have to be provided at a set concentration for sequencing or microarray analysis; sequencing methods report a large but finite total of reads, of which any particular sequence is a proportion. Sometimes, researchers are interested in the relative abundance of different components. Other times, they have to make do with relative abundance to gain insight into the system under study. Whatever the case, data that carry only *relative* information need special treatment.

Awareness is growing [1, 2, 3] but it is not yet widely appreciated that common analysis methods— including correlation—can be very misleading for data carrying only relative information. *Compositional data analysis* [4] (CoDA) is a valid alternative that harks back to Pearson’s observation [5] of *‘spurious correlation’*, i.e., while statistically independent variables *X*, *Y*, and *Z* are not correlated, their ratios *X/Z* and *Y/Z* must be, because of their common divisor. (Note: this differs from the logical fallacy that “correlation implies causation”.)

Proportions, percentages and parts per million are familiar examples of compositional data; the fact that the representation of their components is constrained to sum to a constant (i.e., 1, 100, 10^6^) emphasizes that the data carry only relative information. Note that compositional data do not necessarily have to sum to a constant; what *is* essential is that only the *ratios* of the different components are regarded as informative.

Correlation—Pearson, Spearman or other—leads to meaningless conclusions if applied to compositional data because its value depends on which components are analyzed [4]. Problems with correlation can also be demonstrated geometrically (Figure 1): the bivariate joint distribution of relative abundances says nothing about the distribution of absolute abundances that gave rise to them. Thus, relative data is also problematic for mutual information and other distributional measures of association. To further illustrate how correlation can be misleading we applied it to absolute and relative gene expression data in fission yeast cells deprived of a key nutrient [6].

**Figure 1.**
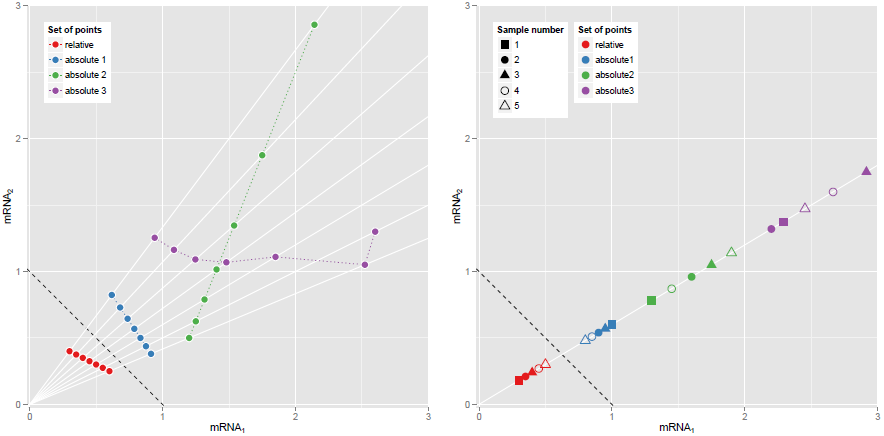
Why correlations between relative abundances tell us absolutely nothing. These plots show two hypothetical mRNAs that are part of a larger total. (a) Seven pairs of relative abundances (mRNA_1_/total, mRNA_2_/total) are shown in red, representing the two mRNAs in seven different experimental conditions.The dotted reference line shows (mRNA_1_ + mRNA_2_)/total = 1.) Rays from origin through the red points show *absolute* abundances that could have given rise to these relative abundances, e.g., the blue, green or purple sets of points (whose Pearson correlations are −1, +1 and 0.0 respectively). (b) Relative abundances that are proportional must come from equivalent absolute abundances. Here the blue, green or purple sets of point pairs have the same proportionality as the pairs of relative abundances in red, though not necessarily the same order or dispersion.

How then can we make sound inferences from relative data? We show how *proportionality* provides a valid alternative to correlation and can be used as the basis of familiar analyses and visualizations. We conclude by putting this analysis strategy in perspective, discussing challenges, caveats and issues for further work, as well as the biological questions raised in this study.

## Results

### Data on absolute mRNA abundance

Our results are based on data from Marguerat *et al.* [6] on the absolute levels of gene expression (i.e., mRNA copies per cell) in fission yeast after cells were deprived of a key nutrient (Figure 2). Unlike many experiments where researchers ensure (or assume) cells produce similar amounts of mRNA across conditions [7], this experiment ensured cells produced very different amounts so as to illustrate the merits of absolute quantification (Figure S14). Total abundance may vary dramatically in other experimental settings—such as in comparing diseased and normal tissues, tissues at different stages of development, or microbial communities in different environments.

**Figure 2.**
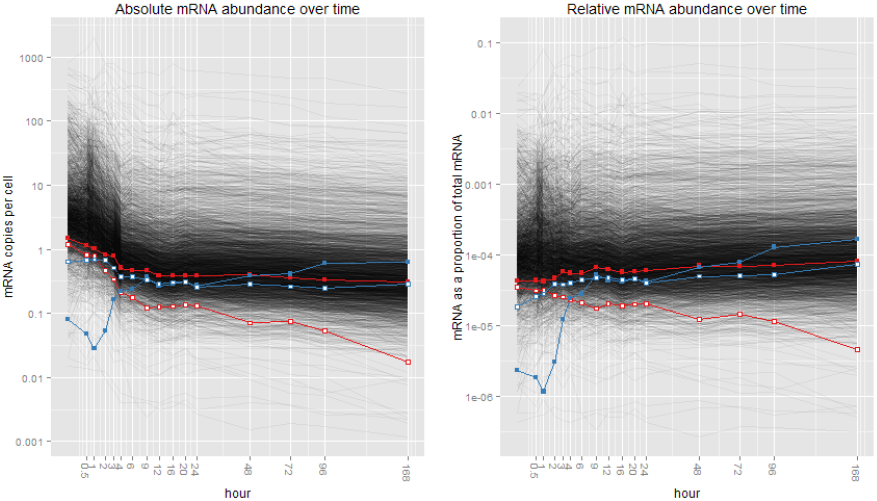
Fission yeast gene expression data of Marguerat *et al.* (a) Absolute and (b) relative abundances of 3031 yeast mRNAs over a 16-point time course. *y*-axes are scaled logarithmically; *x*-axes are on a square-root scale for clarity. Each grey line represents the expression levels of a particular mRNA. The red and blue pairs of mRNAs are discussed later in this paper.

To illustrate the key points of this paper, we worked with positive data only (i.e., we excluded records with any zero or NA values): measurements of 3031 components (i.e., mRNAs) at 16 time points. Furthermore, we applied analysis methods (specifically, correlation) to the absolute abundance data *without* transformation (e.g., taking logarithms) because we believe this approach yields useful insights and simplifies the presentation of the central ideas of this paper (see Section S6.1 and [8]).

### Challenges in interpreting “differential expression”

Before looking at issues with pairs of components, it is important to note that interpreting differences in the relative abundance of a single component can be challenging.

Tests for differential expression are popular for analyzing relative data in bioscience. Much attention has been given to dealing with small numbers of observations and large numbers of tests, but comparatively little to “…the commonly believed, though rarely stated, assumption that the absolute amount of total mRNA in each cell is similar across different cell types or experimental perturbations” [7].

The relationship between the relative and absolute abundance of a component can be understood in terms of fold change over time. When total absolute abundance of mRNA stays constant, fold changes in both absolute and relative abundance of each mRNA are equal. When total absolute abundance varies, fold changes in absolute and relative abundances of each mRNA are no longer equal and can change in *different* directions. Between 0 and 3 hours there were 1399 yeast mRNAs whose absolute abundance *decreased*, and whose relative abundance *increased*. Clearly, mRNAs are being expressed differently, but to describe them as “under- or over-expressed” is too simplistic—here lies the interpretation challenge (see Section S4.4).

### Correlations between relative abundances tell us absolutely nothing

While “differential expression” of relative abundances is challenging to interpret, in the absence of any other information or assumptions, correlation of relative abundances is just wrong. We stress *in the absence of any other information or assumptions* to highlight the common assumption of constant absolute abundance of total mRNA across all experimental conditions. If this assumption holds, and all the mRNAs comprising that total are considered, the relative abundance of each kind of mRNA will be proportional to its absolute abundance, and analyses of correlation or “differential expression” of the relative values will have clear interpretations. The revisitation of this assumption [7] should raise alarm bells about the inferences drawn from many gene expression studies.

Figure 1(a) shows why correlation between relative abundances tells us nothing about the relationship between the absolute abundances that gave rise to them: the perfectly correlated relative abundances could come from *any* set of absolute abundance pairs that lie on the rays from the origin. This many-to-one mapping means that other measures of statistical association (e.g., rank correlations or mutual information) will not tell us anything either when applied to purely relative data.

But is this problem just a theoretical construct? A rare issue? Consider the red mRNA pair in Figure 2: while their *absolute* abundances over time are strongly positively correlated, if someone (inappropriately) used correlation to measure the association between the *relative* abundances of these two mRNAs they would form the opposite view (Figure 3(a)); correlation between the blue mRNA pair in Figure 2 is similarly misleading (Figure S9). What of the other 4.5 million pairs of mRNAs? Figure 3(b) summarizes all discrepancies between correlations of absolute abundance, and correlations of relative abundance, showing clearly that the apparent correlations of relative abundances tell a very different story from those of the absolute data. So how *should* we go about analyzing these relative data?

**Figure 3.**
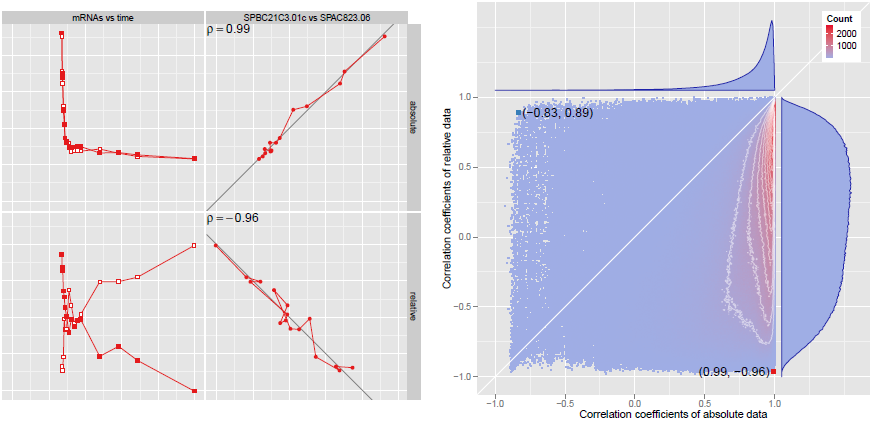
Correlations between relative abundances bear no relationship to the corresponding correlations between absolute abundances. (a) The pair of mRNAs labeled in red in Figure 2, shown on a linear scale. Values have been scaled and translated to have zero mean and unit variance. Upper panels show absolute abundances; the lower show relative abundances. The left panels show mRNA values over time; the right show the value of one mRNA plotted against the other at each time point. The correlation between the relative abundances is almost the complete opposite of that between the absolute abundances of this pair of mRNAs. (b) 2D histogram of the sample correlation coefficient observed for the relative abundances of a given pair of mRNAs, against the correlation observed for the absolute abundances of that same pair, over all pairs. The red and blue points correspond to the red and blue pairs of mRNA in Figure 2. White contour lines are shown at intervals of 100 counts. The top marginal histogram shows that the absolute abundances of most pairs are very strongly correlated. The right marginal histogram shows “the negative bias difficulty”[4].

### Principles for analyzing relative data

CoDA theory provides three principles [4, 9]:

1. Scale invariance: analyses must treat vectors with proportional positive components as representing the same composition (e.g., (2, 3, 4) is equivalent to (20, 30, 40))
2. Subcompositional coherence: inferences about subcompositions (subsets of components) should be consistent, regardless of whether the inference is based on the subcomposition or the full composition.
3. Permutation invariance: the conclusions of analyses must not depend on the order of the components.

Correlation is not subcompositionally coherent: its value depends on which components are considered in the analysis, e.g., if you deplete the most abundant RNAs from a sample [10] and use correlation to measure association between relative abundances, you get different correlations to the undepleted sample (Figure S11).

### Proportionality is meaningful for relative data

Proportionality obeys all three principles for analyzing relative data. If relative abundances *x* and *y* are proportional across experimental conditions *i*, their *absolute* abundances must be in proportion:

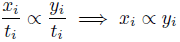

where *t_i_* is the total abundance in condition *i* (Figure 1(b)).

We proposed a “goodness of fit to proportionality” statistic *Φ* to assess the extent to which a pair of random variables (*x, y*) are proportional [11]. *Φ* is related to *logratio variance* [4], var(log(*x/y*)), and is zero when *x* and *y* behave perfectly proportionally. However, when *x* and *y* are not proportional, *Φ* has both a clear geometric interpretation and a meaningful scale, addressing concerns raised about logratio variance [3]: the closer *Φ* is to zero, the stronger the proportionality. We consider “strength” of proportionality (goodness of fit) rather than *testing the hypothesis of proportionality* because it allows us to *compare* relationships between different pairs of mRNAs (Section S5.8).

We calculated *Φ* for the relative abundances of all pairs of mRNAs and compared it to the correlations between their absolute abundances (Figure S19): clearly, the absolute abundances of most mRNA pairs are strongly positively correlated; far fewer are also strongly proportional. Focusing on these strongly proportional mRNAs, we extracted the 424 pairs with *Φ* < 0.05. We graphed the network of relationships between these mRNAs (Figure S28), an approach similar to gene co-expression network[12] or weighted gene co-expression analysis[13] but founded on proportionality and therefore valid for relative data. The network revealed one cluster of 96, and many other smaller clusters of mRNAs behaving proportionally across conditions. Using *Φ* as a dissimilarity measure, we formed heatmaps of the three largest clusters (Figures S30 and S35) similar to the method of Eisen *et al.*[14] but, again, using proportionality not correlation.

## Discussion

This paper does not deny pairwise statistical associations between absolute abundances. What it does say is that once all the absolute information has been removed, only a subset of those associations will reliably endure in the remaining relative data, specifically, associations where values behave proportionally across observations.

### Other approaches to compositional data in the molecular biosciences

Other researchers have recognized the compositional nature of molecular bioscience data, including [15] as discussed in [16]. Strategies have been proposed to ameliorate spurious correlation in the analysis of relative abundances [2, 3]. We contend that there is no way to salvage a coherent interpretation of correlations from relative abundances without additional information or assumptions; our argument is based on Figure 1.

ReBoot [2] attempts to establish a null distribution of correlations against which bootstrapped estimates of correlations can be compared. Aitchison articulates problems with this approach [4, p.56-58]. SparCC [3] injects additional information by assuming the number of different components is large and the true correlation network is sparse. This equates to assuming “that the average correlations [between absolute abundances] are small, rather than requiring that any particular correlation be small” [3, Eq.14]. This means the expected value of the total absolute abundance will be constant (as the sum of many independently distributed amounts). We are concerned with situations where that assumption cannot be made, or where the aim is to describe associations between relative amounts.

### Caution about correlation

We are also keen to raise awareness that correlation (and other statistical methods that assume measurements come from real coordinate space) should not be applied to relative abundances. This is highly relevant to gene coexpression networks [12]. Correlation is at the heart of methods like Weighted Gene Co-expression Network Analysis [13] and heatmap visualization [14]. These methods are potentially misleading if applied to relative data. This concern extends to methods based on mutual information (e.g., relevance networks [17]) since, as Figure 1 shows, the bivariate joint distribution of relative abundances (from which mutual information is estimated) can be quite different from the bivariate joint distribution of the absolute abundances that gave rise to them.

Measures of association produce results regardless of the data they are applied to—it is up to the analyst to ensure that the measures are appropriate to the data. Currently, there are many gene co-expression databases available that provide correlation coefficients for the relative expression levels of different genes, generally from multiple experiments with different experimental conditions (see e.g., [18]). As far as we are aware, none of the database providers explicitly address whether absolute levels of gene expression were constant across experimental conditions. If the answer to this question is “no”, we would not recommend these correlations be used for the reasons demonstrated in this paper. If the answer is “yes” we still advocate caution in applying correlation to absolute abundances for reasons discussed in Section S6.1.

### Results in relation to genome regulation in fission yeast

While the main aim of this study is to present and illustrate principles for analyzing relative abundances, it has also uncovered intriguing biological insight with respect to gene regulation.

The largest cluster of proportionally regulated mRNAs (96 genes, Table S4) was highly enriched for mRNAs down-regulated as part of the core environmental stress response [19], including 66 mRNAs that encode ribosomal proteins, and the remaining mRNAs also associated with roles in protein translation, such as ribosome biogenesis, rRNA processing, tRNA methyltransferases and translation elongation factors. The absolute levels of these mRNAs decrease after removal of nitrogen (Figure S31; [6]). The notable coherence in biological function among the mRNAs in this cluster is higher than typically seen when correlative similarity metrics for clustering are applied (e.g., [19]). These 96 mRNAs show remarkable proportionality to each other over the entire timecourse (Figure S31), and maintain near constant ratios across all conditions (Figure S32). Given the huge energy invested by yeast cells for protein translation (most notably ribosome biogenesis [20, 21], it certainly makes sense for cells to synchronize the expression of relevant genes such that translation is finely tuned to nutritional conditions.

Evidently, numerous ribosomal proteins and RNAs function together in the ribosome, demanding their coordinated expression; more surprisingly, multiple other genes, with diverse functions in translation, show equally pronounced proportional regulation across the timecourse. These findings raise intriguing questions as to the molecular mechanisms underlying this proportional regulation, suggesting sophisticated, coordinated control of numerous mRNAs at both transcriptional and post-transcriptional levels of gene expression.

### Challenges and future work

While proportionality and the *Φ*-statistic provide a valid alternative to correlation for relative data, there are still some challenges in their application. First is the treatment of zeroes, for which there is currently no simple general remedy [22]. Second, and related, is the fact that “many things that we measure and treat as if they are continuous are really discrete count data, even if only at the molecular extremes” [23] and count data is not purely relative—the count pair (1, 2) carries different information than counts of (1000, 2000) even though the relative amounts of the two components are the same. Correspondence analysis [24], or methods based on count distributions (e.g., logistic regression and other generalized linear models) may provide ways forwards.

## Methods

### Reproducing this research

All data and code [25] needed to reproduce the analyses and visualizations set out in this paper are contained in the Supplementary Information, along with additional illustrations and detailed explanations.

### Measuring proportionality

The “goodness of fit to proportionality” statistic *Φ* can be used to assess the extent to which a pair of random variables (*x, y*) are proportional [11]. Aitchison [4] proposed *logratio variance*, var(log(*x/y*)), as a measure of association for variables that carry only relative information. When *x* and *y* are exactly proportional var(log(*x/y*)) = 0, but when *x* and *y* are not exactly proportional, “it is hard to interpret as it lacks a scale. That is, it is unclear what constitutes a large or small value… (does a value of 0.1 indicate strong dependence, weak dependence, or no dependence?)” [3]. Logratio variance can be factored into two more interpretable terms:

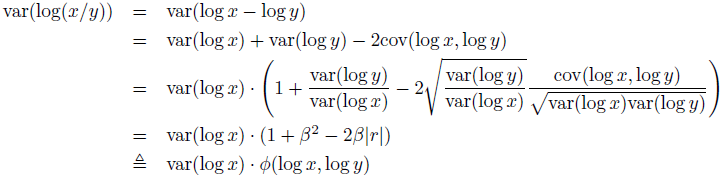

where *β* is the *standardized major axis* estimate [26] of slope of random variables log *y* on log *x*, and *r* the correlation between those variables. The first term, var(log *x*), is solely about the magnitude of variation at play and has nothing to do with *y*. The second term, *Φ*, describes the degree of proportionality between *x* and *y*, and forms the basis of our analysis of the relationships between relative values. Other non-negative functions of *β* and *r* that are zero when *x* and *y* are perfectly proportional could be formed. Section S5 illustrates this, as well as why *Φ* is preferable to an hypothesis testing approach. There is no need to calculate *β* or *r* to assess strength of proportionality; they simply provide a clear geometric interpretation of *Φ*; in practice, one can use the relationship *Φ*(log *x,* log *y*) = var(log(*x/y*))/var(log *x*).

### Centered logratio (clr) representation

We have used *Φ*(log *x,* log *y*) to emphasize the relationship between *Φ* and logratio variance. However to ensure that the *Φ* values for component pair (*i, j*) are on the same scale (i.e., comparable to) the *Φ* values for component pair (*m, n*), it is necessary to use the *centered logratio* (clr) transformation instead of just the logarithm (Section S6). The clr representation of composition **x** = (*x*_1_*,…, x_i_,…, x_D_*) is the logarithm of the components after dividing by the geometric mean of **x**:

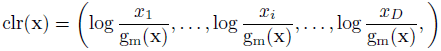

ensuring that the sum of the elements of clr(**x**) is zero. Note that dividing all components in a composition by a constant (i.e., the geometric mean g_m_(**x**)) does not alter the *ratios* of components.

### Using ***Φ*** to form co-expression networks and clustered heatmaps

Gene co-expression networks[12, 13] are generally based on a pairwise distance or dissimilarity matrix which is often a function of correlation and thus not appropriate for relative data. Proportionality is appropriate, but *Φ* does not satisfy the properties of a *distance*—most obviously, it is not symmetric unless *β* = 1:

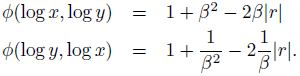

We are most interested in pairs of variables where *β* and *r* are near 1 and want to preserve the link between *Φ*(log *x,* log *y*), *β* and *r*. Hence, our approach to forming a dissimilarity matrix is simply to work with *Φ*(log *x_i_*, log *x_j_*) where *i* < *j*, in effect, the lower triangle of the matrix of *Φ* values between all pairs of components. This symmetrised form of *Φ* was then used to lay out a network of the 145 mRNAs that were involved in 424 pairwise relationships with *Φ* < 0.05 (Section S5.7).

We used the symmetrised form of *Φ* as the basis of the cluster analysis and heatmap expression pattern display described by Eisen *et al.*[14] (Section S5.10).

## Acknowledgments

This research has been supported by CSIRO’s Transformational Biology initiative; by the Spanish Ministry of Education, Culture and Sports under a *Salvador de Madariaga* grant (Ref. PR2011-0290); by the Spanish Ministry of Economy and Competitiveness under the project METRICS Ref. MTM2012-33236; by the *Agència de Gestió d’Ajuts Universitaris i de Recerca of the Generalitat de Catalunya* of the *Generalitat de Catalunya* under project Ref: 2009SGR424; by the UK Medical Research Council; and by a Wellcome Trust Senior Investigator Award (grant #095598/Z/11/Z).

